# Coronavirus-associated molecular mimicry through homology to a SARS-CoV-2 peptide could be leading to susceptibility in patients with HLA-A*02:01 and HLA-A*24:02 serotypes

**DOI:** 10.1101/2021.01.28.428642

**Authors:** Yekbun Adiguzel

## Abstract

This study aims to predict autoimmunity-related pathological mechanisms that possess risk for individuals with specific human leukocyte antigen (HLA) serotypes and shared by certain coronaviruses including SARS-CoV-2, based on homology to a SARS-CoV-2 peptide. With the given aim, 1-) coronavirus-associated sequences, which are homologous to the 15mer SARS-CoV-2 peptide CFLGYFCTCYFGLFC, are obtained. 2-) Human peptides that have at least 7 residue matches with those coronavirus sequences, and the SARS-CoV-2 15mer, are found. 3-) Epitope pairs, which are sourced by those aligned coronavirus and human sequences are identified. 4-) Epitope pairs that are predicted to bind strongly not only to the same HLA allele with each other but also to the same HLA allele as those of the respective alignment of the SARS-CoV-2 peptide are selected. Following are the identified proteins or peptides (with HLA-A*02:01 or HLA-A*24:02 epitopes), as described in 1-to-4: Immunoglobulin heavy chain junction regions, CRB1 isoform I precursor, slit homolog 2 protein, hCG1995581, hCG2028737, phospholipid phosphatase-related protein type 2. Among those, CRB1 isoform I precursor sequence with the predicted HLA-A*24:02 epitope aligns with the highest number of different sequences. Results imply autoimmunity risk in COVID-19 patients with HLA-A*02:01 and HLA-A*24:02 serotypes, through molecular mimicry, as a shared pathogenicity risk that can be prevalent upon getting infected with certain coronaviruses. These can pave way to improved risk groups’ assessment and autoimmunity treatment options, for COVID-19 and its associated diseases. Also, the approach in this study can be used to predict prospective pathologies of the transmissible variants in susceptible humans.

## Introduction

Upon infection, presence of similar sequences to the pathogens’ proteins in the human proteome can be a potential risk of causing autoimmune response. This molecular mimicry is among the preliminary conditions. Kerkar and Vergani [1] mentioned molecular mimicry, sequence similarities of pathogenic organisms with autoantigens, as a possible mechanism of autoimmunity, and is conceptually reinforced through the discoveries of de novo autoimmune hepatitis associated with certain viral infections. In 90s, a sequence similarity search revealed 70 % overlap of a 10mer within the V3-loop of the envelope glycoprotein gp120 of HIV-1 isolates, with the collagen-like region of the human complement component C1q-A [2]. Follow-up studies by the same group addressed the complications that would be caused by vaccines, which are based on gp120 of HIV-1 [3]. Later, their findings on the presence of antibodies that are reactive for the peptide in the V3-loop of HIV-1 in the healthy individuals [4], and the complementarity of the antibodies for V3-loop of HIV-1 and IgG of human [5] were reported. Presence of homologies between human proteins and virus proteins, and the examination of the pathways of those homologous peptides, drew attention to the pathologies of viral infections that are related to the immune response [6]. Even therapies targeting such complexifications were suggested. In correlation with these, Kanduc and Shoenfeld [7] considered the intrinsic hazards in the vaccines that are based on such pathogen sequences. The authors later extended their earlier findings, and the theoretical implications of those [8,9]. Similarities among the SARS-CoV-spike glycoprotein sequences and those of the human surfactant protein and the related proteins were revealed by the authors, through a similar methodology [10]. Specifically, 13 of 24 shared pentapeptides were found to be present in 52 SARS-CoV-derived immunoreactive epitopes. In addition, heptapeptide-sharing among pathogens, which are also containing SARS-CoV-2, and the non-human primates, was observed and it revealed that high level of peptide-sharing is somewhat unique to human [11]. The findings of their studies also led the authors to suggest “aged mice” for testing of vaccines that are based on spike glycoproteins of SARS-CoV-2 [12].

Relationship of COVID-19 and immunity is complex [13,14,15,16,17,18], it can involve autoimmune reactions through molecular mimicry. Woodruff and co-workers [19] reported that critically ill patients of COVID-19 “displayed hallmarks of extrafollicular B cell activation as previously described in autoimmune settings.” Bastard and co-workers [20] reported autoantibodies against type I IFNs in people with serious COVID-19 disease. Rodríguez and co-workers [21] reviewed autoinflammatory and autoimmune conditions in COVID-19. Accordingly, antiphospholipid syndrome, autoimmune cytopenia, Guillain-Barré syndrome and Kawasaki disease are known to be reported in COVID-19 patients [21]. A more recent review [22] is present as well. Regarding molecular mimicry, Cappello and co-workers [23] hypothesized that SARS-CoV-2 may be generating autoimmunity through molecular mimicry, induced by stress. Rodríguez and co-workers [21] mentioned the molecular mimicry and bystander activation as the mechanisms that can link COVID-19 to autoimmunity. In relation, Lucchese and Flöel [24] reported three 6mers in the human brainstem respiratory pacemaker proteins that are present in the SARS-CoV-2 proteome. The authors also reported molecular mimicry with SARS-CoV-2 and heat shock proteins 90 and 60 of human, which are known to be associated with Guillain-Barré syndrome and other autoimmune diseases [25]. Importantly, those shared peptides are part of those epitopes that were experimentally shown to be immunoreactive. Also, Kanduc and Shoenfeld [10] reported 5mers of human surfactant protein to be present in the SARS-CoV-2 proteome. Angileri and co-workers [26] reported a 7mer of human Odorant Receptor 7D4, a 6mer of human Poly ADP-Ribose Polymerase Family Member 9, and a 7mer of Solute Carrier Family 12 Member 6, which are present in the putative epitopes of SARS-CoV-2. There are also human proteins that have strong immune cross-reactions with the spike protein antibody of SARS-CoV-2 [27], which can be suggestive of molecular mimicry-based autoimmunity in susceptible individuals [21]. In relation, we looked for autoimmunity related pathological mechanisms that are common to certain coronaviruses, including SARS-CoV-2, by means of a selected sequence (CFLGYFCTCYFGLFC), which is obtained through our ongoing study [28] involving tblastx search of SARS-CoV-2 and plasmodium species that cause malaria in human [29,30].

## Materials and Methods

The CFLGYFCTCYFGLFC sequence was obtained by performing blastx [31] at NCBI [32], between the reference genome of query input SARS-CoV-2 (NC_045512.2) and *P. vivax* (taxid:5855) [28]. It was the aligned query sequence in the tblastx output that revealed the top identity between the query and subject, which was afterwards utilized as input for NCBI blastp search, by limiting the search to SARS-CoV-2 (taxid:2697049), to ensure that the sequence is expressed [28]. Here, in this study, blast of the sequence is performed at Uniprot with threshold 10 (*performed on 5 August 2020*). The associated coronaviruses with homologous sequences are selected from the results. Then, blastp of the SARS-CoV-2 sequence and the coronavirus sequences that are homologous, are performed separately, by limiting the searches to *H. sapiens* (taxid:9606). The query-subject sequence pairs that have at least 7 residue matches are found in those results. Within those results, the ones with the same names and sequence IDs as those in the respective results of SARS-CoV-2 blastp search, are identified. This is followed by major histocompatibility complex (MHC) class I binding predictions for those identified query-subject sequence pairs. This is done to find the coronavirus and human sequences that are homologous, and which are predicted to bind strongly to the same human leukocyte antigen (HLA) alleles as that with the SARS-CoV-2 sequence. To do that, binding affinities to the MHC class I (*MHC class I genes are HLA-A, -B, and -C genes* [33]) proteins are predicted with the use of a tool that integrates NetMHC 4.0 [34,35], NetMHCpan 4.1 [36], and PickPocket 1.1 [37]. That tool is NetMHCcons 1.1 [38]. Predictions are performed for 8-15mers, with default parameters, and by performing the predictions for 12 MHC supertype representatives. The 12 MHC supertype representatives are HLA-A*01:01 (A1), HLA-A*02:01 (A2), HLA-A*03:01 (A3), HLA-A*24:02 (A24), HLA-A*26:01 (A26), HLA-B*07:02 (B7), HLA-B*08:01 (B8), HLA-B*27:05 (B27), HLA-B*39:01 (B39), HLA-B*40:01 (B44), HLA-B*58:01 (B58), HLA-B*15:01 (B62). NetCTLpan 1.1 [39] is utilized as well, similarly, for the prediction of the epitopes of cytotoxic T lymphocyte (CTL), as 8-11mers. Within the prediction results, epitopes with at least 5 residue matches and strong binding affinities to the same MHC supertype representative are considered for possible risk of autoimmunity-related pathological mechanisms that are common to SARS-CoV-2 and associated coronaviruses, based on similarity to the selected short SARS-CoV-2 sequence. Epitope-pairs with the highest number of residue-matches are displayed.

*Summary information* - Summary information about the identified proteins (or peptides in case of immunoglobulin heavy chain junction regions) is collected by searching the sequence ID of the aligned subject sequence in the blastp results at Entrez (NCBI), and then searching the encoding gene ID, which is indicated at the UniProt (https://www.uniprot.org) [40], to retrieve the UniProtKB number. Information on the associated diseases is obtained from the human gene database GeneCards (https://www.genecards.org) [41], wherever readily available.

## Results

The query peptide with the sequence CFLGYFCTCYFGLFC in the single letter code representation is originally obtained through tblastx search (see s2 file of [28]). Peptide with that sequence is present in the SARS-CoV-2 proteome (see s6 file of [28]), as part of non-structural protein 6 (nsp6), which is cleaved from the replicase polyprotein 1a. Blast search of the sequence is performed (*on 5 August 2020*) at Uniprot, with threshold 10 (S1, *all Sfiles are deposited at shorturl.at/rswS6*). The associated coronaviruses are retrieved from that search. They are presented in Table 1, along with their related sequences.

**Table 1.**
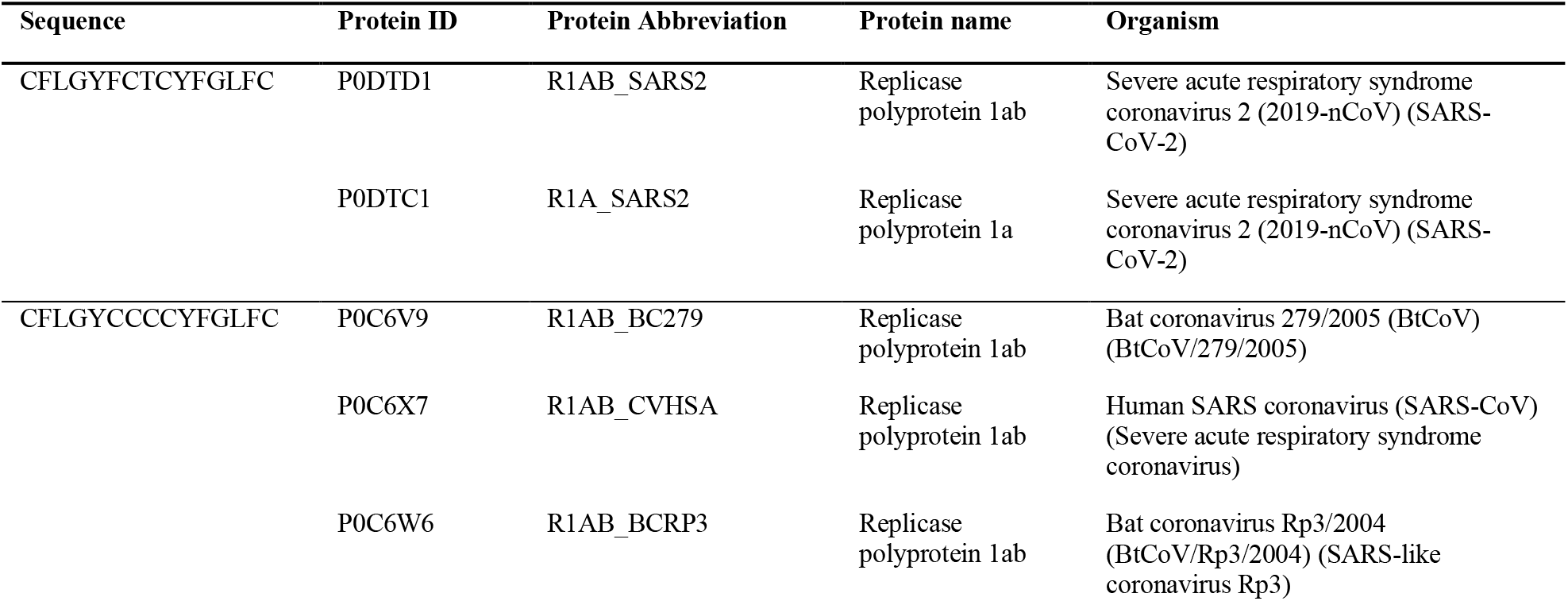
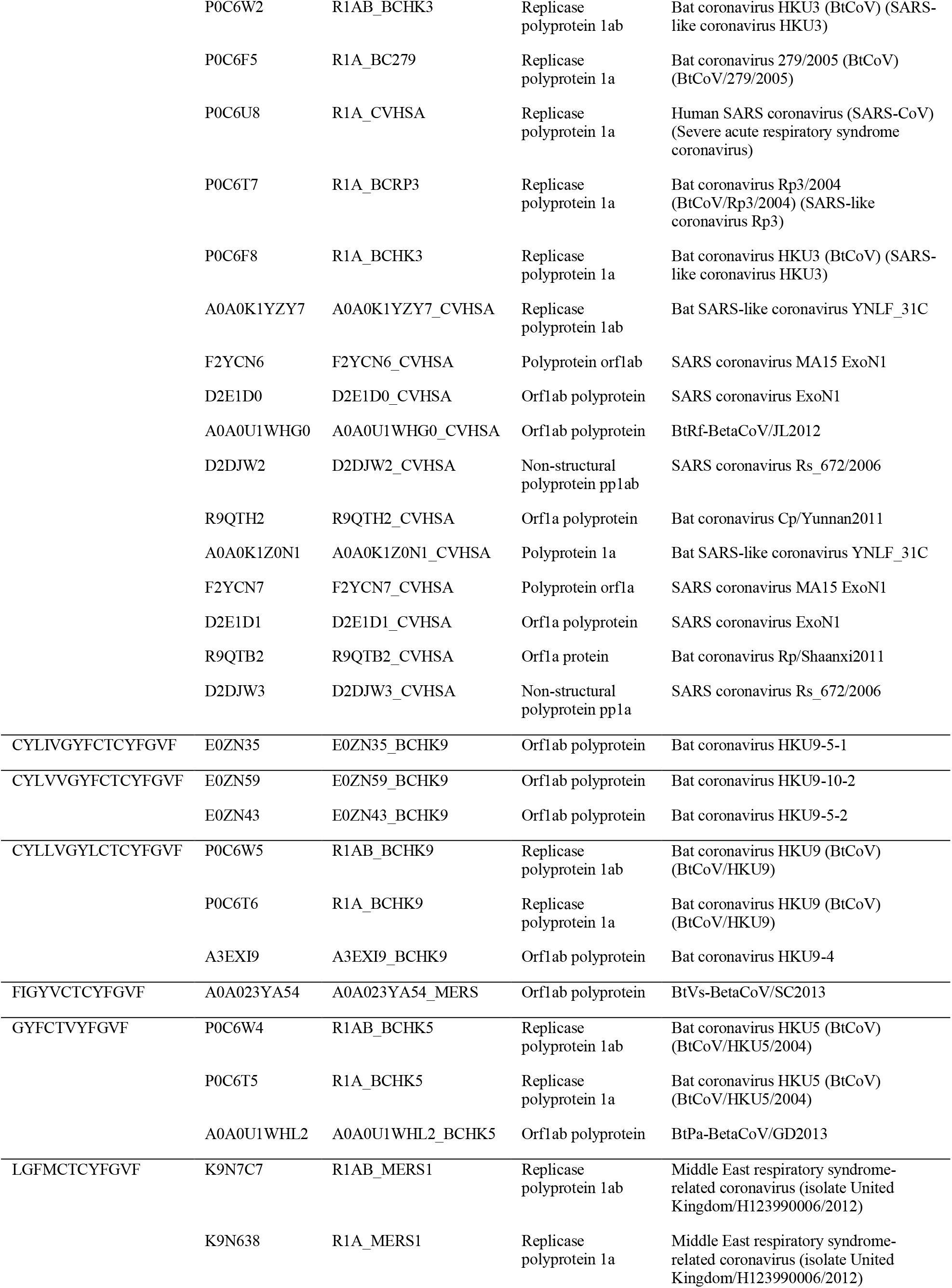
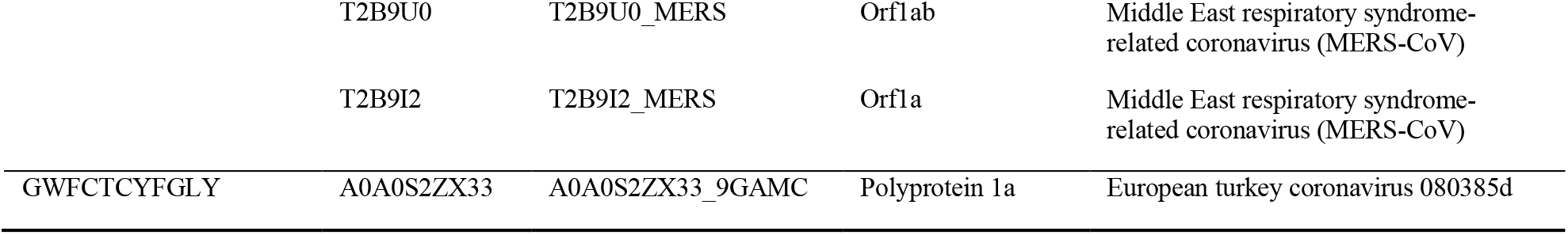
Associated coronaviruses with their sequences that are obtained from the blast search results of the CFLGYFCTCYFGLFC, performed at Uniprot with threshold 10.

Blastp search results of the sequences that are displayed in the first column of Table 1 are obtained by limiting the search to *H. sapiens* (S2-10). The query-subject sequence pairs in those results, which have at least 7 residue matches, are found (S11, *file displays the respective subject sequences and the proteins containing them*). Among those, the ones with the same protein name and sequence IDs as those in the respective results of SARS-CoV-2 blastp, are identified (Table 2). The LGFMCTCYFGVF sequence displayed at Table 1 do not have any alignment with the same protein name and sequence ID as those that are aligned with the SARS-CoV-2 sequence. The GWFCTCYFGLY sequence displayed at Table 1 do not have any alignment with the same protein name and sequence ID neither as those aligned with the SARS-CoV-2 sequence nor as the other coronaviruses. Yet, we looked for at least 7 residue matches, indicating that the outcome could be different with 6 residue matches, or so.

**Table 2.**
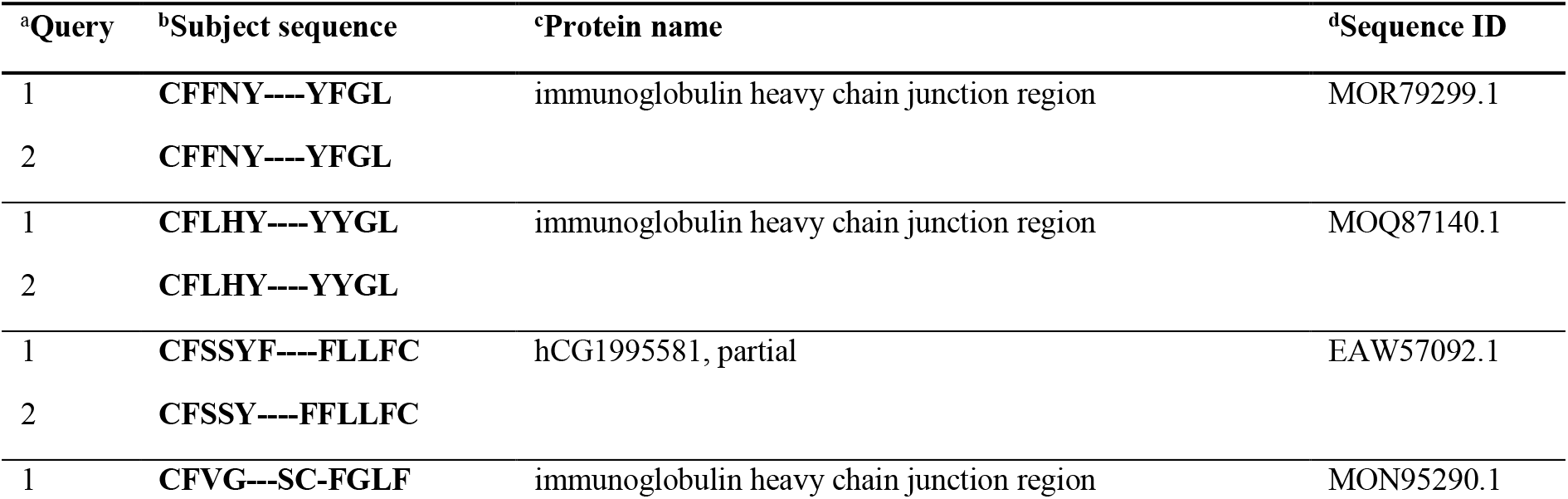
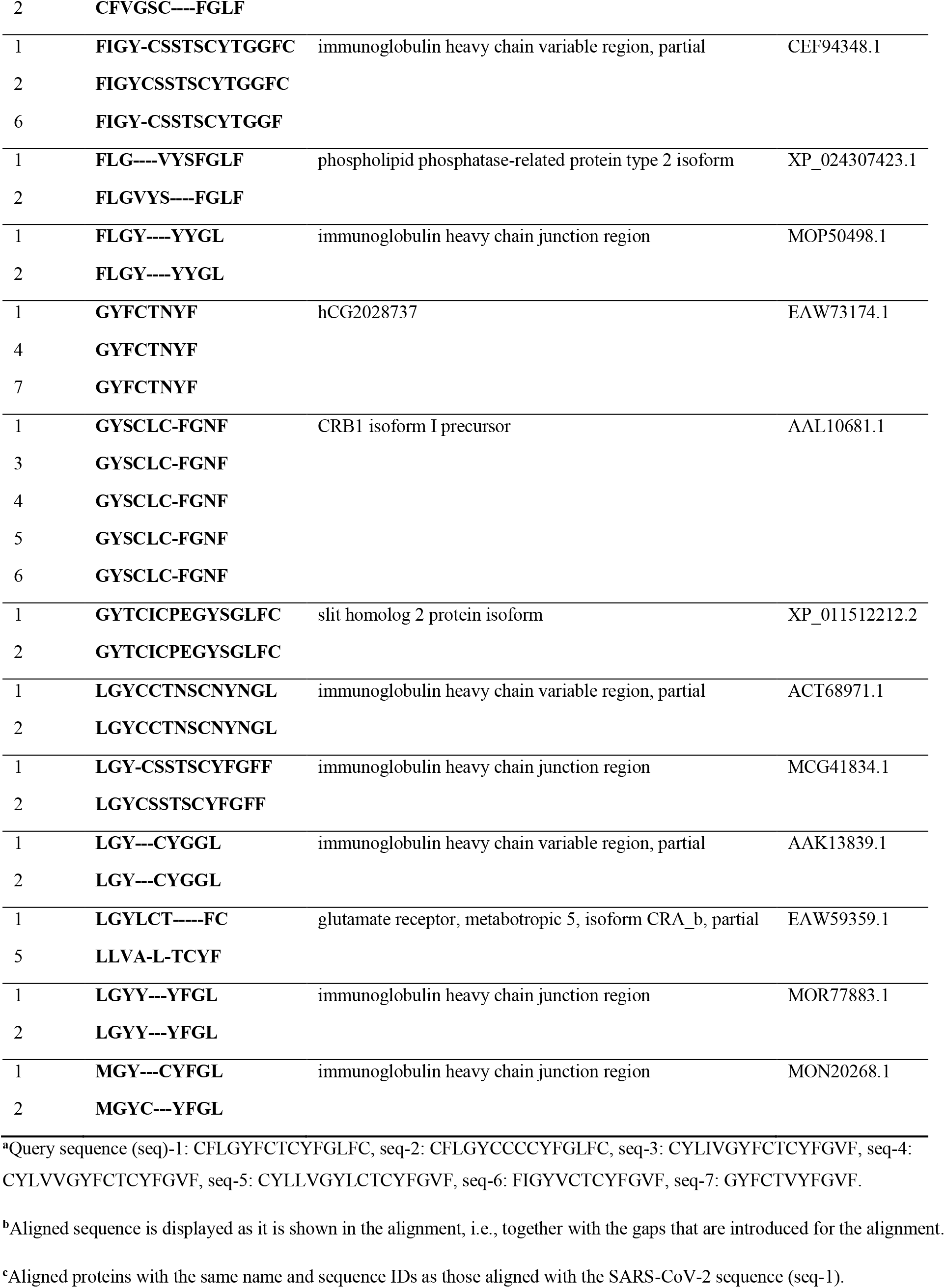
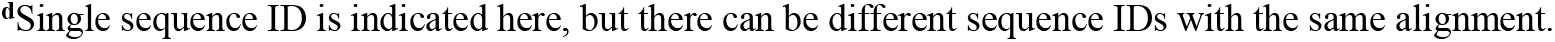
Human proteins and their subject sequences, which are aligned with the coronavirus sequences that are homologous with the 15mer SARS-CoV-2 sequence (query number 1, seq-1). Aligned subject sequences that are shown in the second column contain at least 7 residues that are matching with the query sequences, which are indicated by a designating-number in the first column. Descriptions of those numbers are at the end of the table. Aligned sequences in the second column are displayed together with the gaps that are introduced to the sequence for the alignment.

It is observed at Table 2 that there are 24 different alignments of 16 human protein sequences with the coronavirus sequences. All the listed human proteins in Table 2 are aligning with seq-1, which is a SARS-CoV-2 sequence, and with at least one other coronavirus sequence that is homologous to seq-1. MHC supertype representative-binding predictions are performed for (both the query and subject) sequences that are displayed at Table 2. Table 3 presents the MHC supertype representative-binding predictions for the homologous coronavirus (query) and human (subject) sequence pairs, which are predicted to bind strongly to the same HLA alleles, not only with each other, but also with those of the respective SARS-CoV-2 (query) and human (subject) sequences. Immunoglobulin heavy chain junction regions with sequence IDs MOR79299.1, MOR77883.1, and MON20268.1; immunoglobulin heavy chain variable regions with sequence IDs ACT68971.1 and AAK13839.1; and metabotropic glutamate receptor 5 isoform with sequence ID EAW59359.1 are displayed in Table 2, but not in Table 3. The query-subject sequence pairs with those protein sequences are not predicted to bind strongly to the same HLA alleles. On the other hand, immunoglobulin heavy chain variable region with the sequence ID CEF94348.1 is also absent in Table 3 because the query-subject sequence pairs of those other than that of the SARS-CoV-2 sequence are not predicted there, to be binding strongly to the same HLA alleles.

**Table 3.**
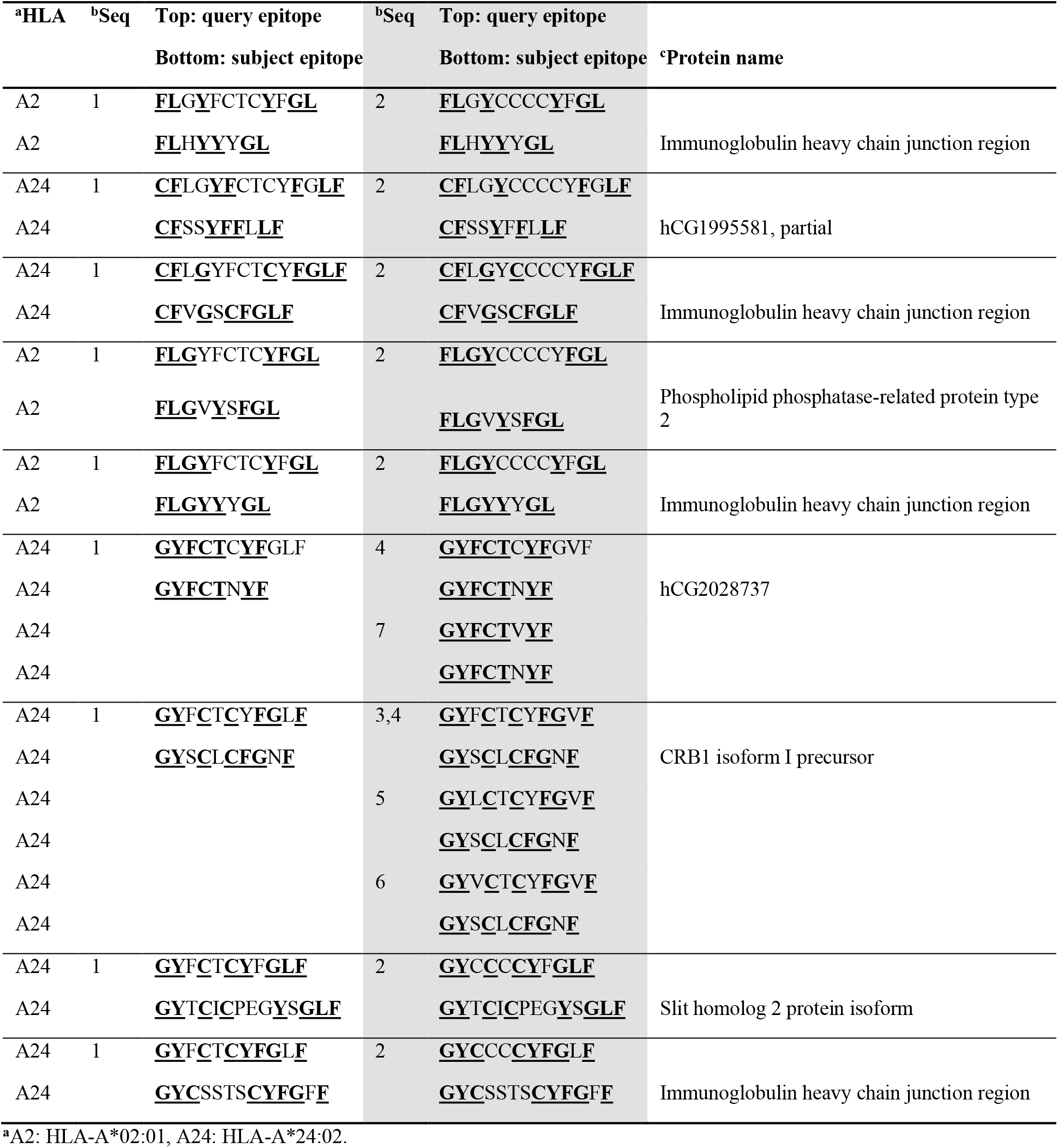
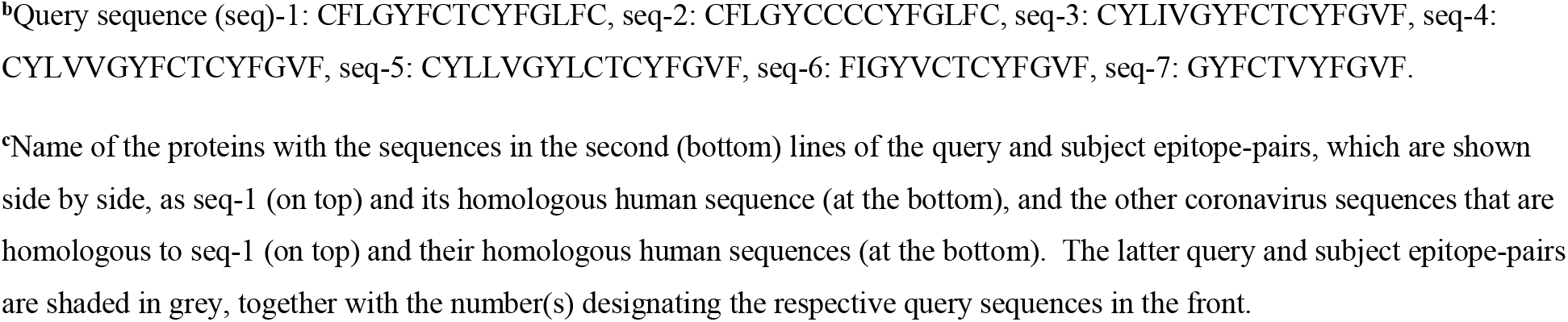
Human proteins and their (subject) sequences, which are aligned with the coronavirus (query) sequences that are not only homologous to the SARS-CoV-2 15mer (seq-1) but also predicted to bind strongly to the same HLA alleles as that of the SARS-CoV-2 (query) and human (subject) epitope-pairs. These epitope-pairs are shown in consecutive rows and each epitope-pair have at least 5 matching-residues. Residues in these epitope-pairs are written bold if they are among the matching-residues in the original alignments. Those residues are further underlined if they are still present as matching-residues in the predicted epitope-pairs. HLA allele is indicated in the first column. Accordingly, the same HLA allele is displayed in case of the coronavirus (query) and human (subject) epitope-pairs. Epitopes predicted by NetMHCcons are displayed here. Epitope-pairs that are sourced by the alignments of the SARS-CoV-2 15mer (seq-1) are shown in the third column. In the second column, 1 indicates the SARS-CoV-2 (query) sequence-number and it is written at the rows, which correspond to the same rows as the SARS-CoV-2 (query) epitopes of the epitope-pairs at the third column. Epitope-pairs that are sourced by the alignments of the other coronavirus sequences are shown in the fifth column. The numbers in the fourth column indicates the other coronaviruses’ (queries’) sequence-numbers and they are written at the rows, which correspond to the same rows as the respective coronaviruses’ (queries’) epitopes of the epitope-pairs at the fifth column. Names of the human proteins with the aligned subject sequences are displayed at the last column.

The predictions displayed in Table 3 belong to NetMHCcons, but predictions are performed with NetCTLpan as well. Its results were similar for those that have high affinity to the HLA-A*24:02 allele, according to the NetMHCcons. Exceptions are the predictions for the alignments of seq-2 (CFLGYCCCCYFGLFC) with the hCG1995581 (sequence ID EAW57092.1); predictions for the alignments of both seq-1 and seq-2, with the slit homolog 2 protein (sequence ID XP_011512212.2); and the alignments of seq-2 with one of the immunoglobulin heavy chain junction regions (sequence ID MCG41834.1).

It is observed in Table 3 that the predicted epitope-pairs, which include peptides that are sourced by two immunoglobulin heavy chain junction regions (sequence IDs MOQ87140.1 and MOP50498.1) and phospholipid phosphatase-related protein type 2 isoform (sequence ID XP_024307423.1), are predicted to bind strongly to the HLA-A*02:01 (A2) allele. On the other hand, those that include peptides sourced by the hCG1995581 (sequence ID EAW57092.1); two other immunoglobulin heavy chain junction regions (sequence IDs MON95290.1 and MCG41834.1); hCG2028737 (sequence ID EAW73174.1); CRB1 isoform I precursor (sequence ID AAL10681.1); and slit homolog 2 protein isoform (sequence ID XP_011512212.2), bind strongly to the HLA-A*24:02 (A24) allele. Peptides of most of these proteins that overlap with seq-1, the SARS-CoV-2 peptide’s sequence, overlap also with seq-2, the coronavirus peptide with the sequence CFLGYCCCCYFGLFC. This is expected since seq-2 is closer in sequence to the SARS-CoV-2 peptide (seq-1), compared to the others (Figure 1). Yet, the CRB1 isoform I precursor is the protein with an antigenic peptide with similarity to the highest number of different coronavirus sequences (seq-3 to −6), in addition to the SARS-CoV-2 sequence (seq-1) (Table 3).

**Fig. 1.**
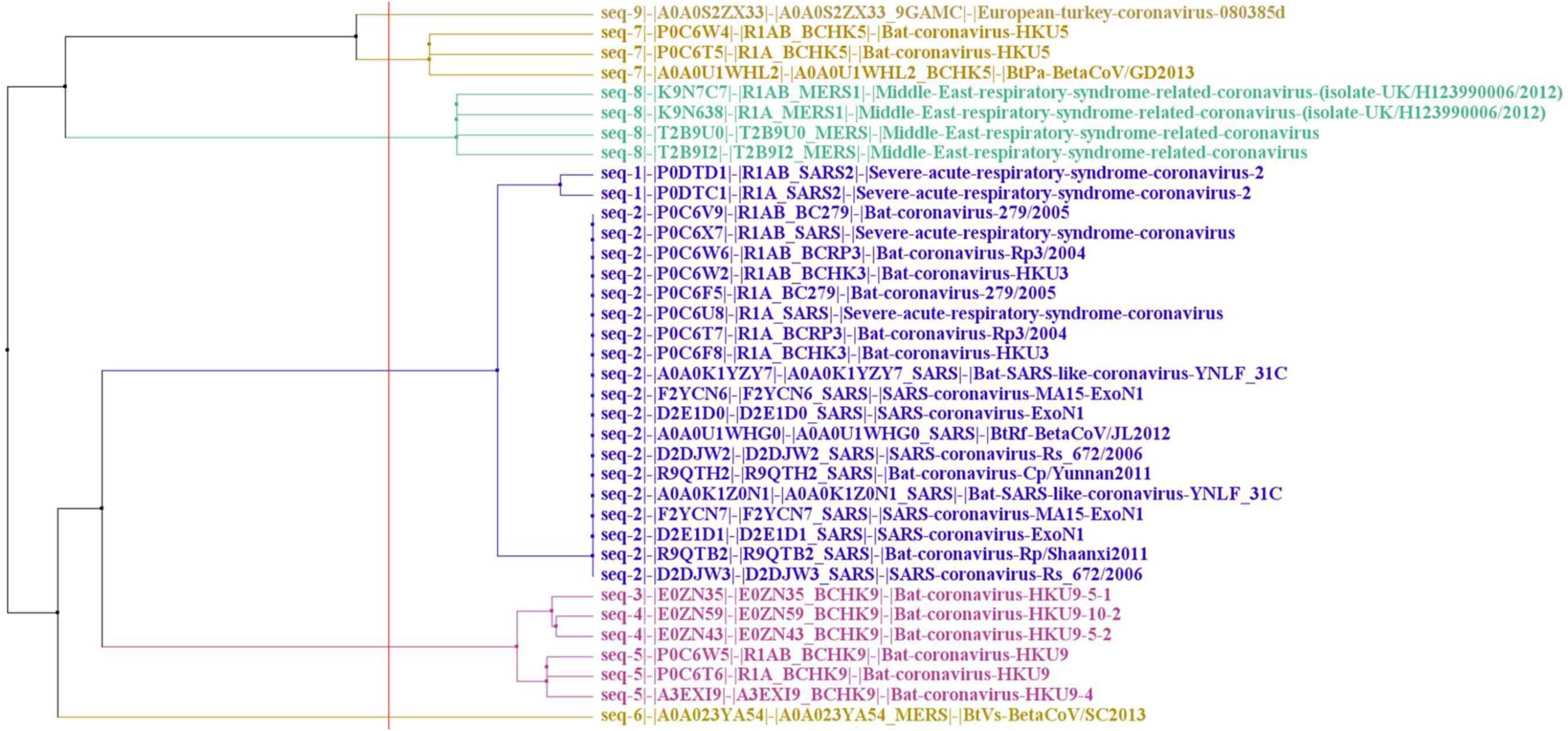
Tree calculated with the alignment of 8 coronavirus sequences homologous to the SARS-CoV-2 sequence CFLGYFCTCYFGLFC (seq-1), and the SARS-CoV-2 sequence (seq-1) itself.^11^ Sequences are obtained by performing Blast of the SARS-CoV-2 sequence at Uniprot, with threshold 10 (S1, *performed on 5 August 2020*). The tree is calculated with Jalview (version 2.11.1.3) [42], Blosum 62, average distance.

Summary information of all the proteins (or peptides) in Table 3:

### Immunoglobulin heavy chain junction region

– 1 alignment in case of SARS-CoV-2 query sequence
– NCBI sequence ID MOQ87140.1
– 22 (aa) (*NCBI*)

### hCG1995581

– 1 alignment in case of SARS-CoV-2 query sequence
– NCBI sequence ID EAW57092.1

### Immunoglobulin heavy chain junction region

– 2 alignments in case of SARS-CoV-2 query sequence, which are the same, except that they are at 2 different subject-sequence regions
– NCBI sequence IDs MON77051.1 and MON95290.1
– 19 aa (*NCBI*)

### Phospholipid phosphatase-related protein type 2

– 12 alignments in case of SARS-CoV-2 query sequence, which are the same, except that they are at 7 different subject-sequence regions
– UniProtKB - Q96GM1 (PLPR2_HUMAN)
– Gene PLPPR2
– Has binary interactions with 21 proteins (*UniProt*)

### Immunoglobulin heavy chain junction region

– 1 alignment in case of SARS-CoV-2 query sequence
– NCBI sequence ID MOP50498.1
– 26 aa (*NCBI*)

### hCG2028737

– 1 alignment in case of SARS-CoV-2 query sequence
– NCBI sequence ID EAW73174.1

### Protein crumbs homolog 1

– 17 alignments in case of SARS-CoV-2 query sequence, which are the same, except that they are at 13 different subject-sequence regions
– UniProtKB - P82279 (CRUM1_HUMAN)
– Gene CRB1
– Takes role in photoreceptor morphogenesis and may maintain cellular adhesion and polarization (*UniProt*)
– Has binary interactions with PATJ [Q8NI35] (*UniProt*)
– Associated diseases involve Leber Congenital Amaurosis 8 and Retinitis Pigmentosa 12 (*GeneCards*)

### Slit homolog 2 protein

– 16 alignments in case of SARS-CoV-2 query sequence, which are the same, except that they are at 12 different subject-sequence regions
– Neurogenic extracellular slit protein Slit2 is also the name that is given at the blastp results
– UniProtKB - O94813 (SLIT2_HUMAN)
Gene SLIT2
– Believed to function as a molecular guidance cue in cellular migration and it takes roles in axonal guidance and development of the parts of the neural system (*UniProt*)
– Has binary interactions with itself and ROBO1 [Q9Y6N7], which is its receptor (*UniProt*)
– Associated diseases involve Cakut and Crohn’s Colitis (*GeneCards*)

### Immunoglobulin heavy chain junction region

– 1 alignment in case of SARS-CoV-2 query sequence
– NCBI sequence ID MCG41834.1
– 21 aa (*NCBI*)

## Discussion

Bianchi and co-workers [43] indicated earlier that studies involving peptide elution confirmed the presentation of transmembrane helices by the MHC class I molecules. In accordance, CFLGYFCTCYFGLFC is likely a transmembrane region, and is predicted to bind strongly to certain HLA alleles, like its homologous sequences in the human proteome [44]. Here, the subset of those that are common to the other coronaviruses is found (Table 3). Some associated diseases of the proteins with the aligned peptides as potential epitopes are mentioned in the’ summary information’. In relation to this study, Lyons-Weiler [45] identified immunogenic epitopes in the proteome of SARS-CoV-2. They compared those epitopes with the proteome of human. More than 1/3 of the immunogenic peptides were found to be homologous to the proteins that are significant in the adaptive immune system. Kanduc [46] suggested that there is an extensive range of health disorders, related to probable autoimmune reactions against the peptides of the human proteome that are homologous to the immunogenic peptides of SARS-CoV-2. It was indicated in another study [47] that cerebrospinal fluids from the patients of COVID-19 were suggestive of autoimmunity. In line with those, results presented here suggest molecular mimicry-based autoimmunity risk in COVID-19 patients with HLA-A*02:01 and HLA-A*24:02 serotypes, as a shared risk to being exposed to certain coronaviruses. Immune responses to the peptides and proteins with similar sequences and strong binding affinities to the same HLA allele can lead to autoimmune reaction [48,49,50,51,52,53,54]. Yet, Trost and co-workers [51] pointed at dramatically high number of shared heptapeptides (7mers) between bacterial and human proteomes. In relation, Amela and co-workers [55] also demonstrated that most of the pathogen proteins that can cause antibody generation by the host immune system are not homologous to the human proteins, and *vice versa*. Distinguishing self from non-self was suggested to be contributing this observation [56]. There need to be genetic, physiological, and environmental [57] variations in action. Additional proofs, although some are challenging, and not free from concerns, are required in addition to (peptide-)similarity. Autoimmune reactions were shown to be developed in the animal models under such conditions (e.g. [58,59]).

Involvement of evolutionary processes [60] is a related concern, not only to these discussions but also to this study and its results. Vaccine targets are already studied for COVID-19 [61], but cross-reactivity risk in the adjuvant-vaccine for the individuals with genetic susceptibility should also be considered [62], together with the discussed considerations in terms of the autoimmunity risk, in general. Similar studies as this one, and as those mentioned here [7,8,9,10,11,12,45,46], and relevant studies [63], all need to consider the possible variations in the genetic makeup as well.

As stated in section 3, the results section, CRB1 isoform I precursor is the protein with an antigenic peptide with similarity to the highest number of different coronavirus sequences (seq-3 to -6), in addition to the SARS-CoV-2 sequence (seq-1) (Table 3). Accordingly, the epitope sourced by the CRB1 isoform I precursor is a strong candidate of a common autoimmune reaction risk source, in case of individuals with the HLA-A*24:02 serotype, upon getting infected by SARS-CoV-2 or the other coronaviruses. In support, CRB1 related coronavirus pathogenicity is previously suggested [64,65]. It is suggested here further that possible CRB1 related coronavirus pathogenicity is additionally fostered by autoimmunity risk in susceptible individuals. However, seq-2, which is including SARS-CoV-1 among the source species, is not among those different coronavirus sequences (seq-3 to -6). This result is obtained through homology to one SARS-CoV-2 sequence (seq-1), which is limited. Yet, this result is in line with related studies. De Maio and co-workers [64] pointed at the importance of Envelope (E) protein of SARS-CoV-2, which is significantly different from SARS-CoV, and bats and pangolin coronaviruses. E protein was reviewed by Schoeman and Fielding [66], in terms of its possible relation with the immunopathology of the disease. E protein binds with the tight-junction associated PALS1 protein, leading to redistribution to the ER-Golgi intermediate compartment and disruption of tight junction, and epithelial barrier damage [66]. De Maio and co-workers [64] suggested that the strengthened binding of E protein with PALS1, which is supported by their later studies [67], can increase epithelial barrier weakening, inflammatory events, and tissue remodeling. The latter, tissue remodeling, is also mentioned by Ziegler and co-workers, as de-differentiation [68].

CRB1 is natural ligand of PALS1 PDZ domain [67]. Autoimmune response to CRB1 through molecular mimicry can reduce the availability of natural ligand CRB1 of PALS1 and destabilize PALS1, or alter its function, in addition to the mechanisms that are discussed above. It can also be mentioned regarding the possible autoimmune response related pathologies in susceptible individuals with COVID-19 that CRB1 is among the top affiliated genes of many diseases that are related to the rare disease Late-Onset Retinal Degeneration, as seen at MalaCards [69]. The site (https://www.malacards.org/card/late_onset_retinal_degeneration?showAll=True) displays through text searches within MalaCards or GeneCards Suite gene sharing. CRB1 is also among the important genes related to retinis pigmentosa [70]. In relation, our findings suggest possible autoimmune reaction development against CRB1 protein in susceptible individuals, upon getting infected with SARS-Co-V-2 or MERS (BtVs-BetaCoV/SC2013). Interestingly, this is not the case for seq-2, which is including SARS-CoV-1 among the source species, while bat coronavirus (HKU9) sequences (seq-3 to -5) have the same property. However, high affinity of those bat coronavirus (HKU9) sequences (seq-3 to -5) to the same HLA allele as human CRB1 peptide is not necessarily suggestive of a probable autoimmune reaction risk in susceptible animals. Yet, this approach can be used to predict prospective pathologies of the transmissible variants in susceptible humans.

As mentioned, it is observed here that the epitope sourced by the CRB1 isoform I precursor is the candidate of a common coronavirus-sourced autoimmune reaction risk in the individuals with the HLA-A*24:02 serotype, upon getting infected. Warren and Birol [71] also identified the same HLA allele in predictions from the transcriptome sequences of bronchoalveolar lavage fluid of 4 among 5 patients from China, with COVID-19. Yet, different from here, the allele was reported not to be (recognized as) a SARS-risk-factor. Findings of this work will be investigated further in the follow-up studies.

## Conclusion

We investigated autoimmunity related, shared pathological mechanisms of coronaviruses, through a 15mer SARs-CoV-2 peptide with the sequence CFLGYFCTCYFGLFC. Accordingly, coronavirus sequences homologous to the SARS-CoV-2 peptide are initially obtained. Afterwards, those homologous coronavirus sequences that are aligning with the human proteins with at least 7 residue matches are attained. They were also expected to be common to the similar alignments of the SARS-CoV-2 peptide with the human proteins. So, coronavirus (query) and human (subject) epitope-pairs are identified. Those epitope-pairs were not only predicted to bind strongly to the same HLA alleles with each other but were also predicted to bind strongly to the same HLA alleles as the respective peptides of the SARS-CoV-2 15mer, and the human protein(s) aligning with it. Immunoglobulin heavy chain junction regions, phospholipid phosphatase-related protein type 2, CRB1 isoform I precursor, and slit homolog 2 protein are among the proteins with those predicted epitopes. They can be related to pathological conditions in susceptible patients. Results suggest autoimmune reaction risk related to the CRB1 protein, in individuals with the HLA-A*24:02 serotypes, upon getting infected with SARS-Co-V-2 or MERS (BtVs-BetaCoV/SC2013). This is suggested also to be the case in bat coronavirus (HKU9). High affinity of bat coronavirus (HKU9) sequences to the same HLA allele as human CRB1 peptide is not necessarily suggestive of a probable autoimmune reaction risk in susceptible animals. Yet, this approach can be used to predict prospective pathologies of the transmissible variants in susceptible humans. Overall, the results here infer molecular mimicry-based autoimmunity risk in individuals with HLA-A*02:01 and HLA-A*24:02 serotypes, in general, upon getting infected with certain coronaviruses, including SARS-CoV-2. These findings can pave the way to clinical studies for autoimmunity treatment options to be used in COVID-19 and associated diseases; identification of individuals with specific HLA alleles as disease risk groups; and consideration of genetic susceptibilities in vaccine studies; and can be further nourished by the identification of novel alleles [72,73].

## Supporting information

Supplementary files available at shorturl.at/rswS6

## Acknowledgments

Ecology and Evolutionary Biology Society of Turkey is acknowledged.

## Authorship

YA performed conception and design of the work; acquisition, analysis, and interpretation of data for the work; drafting and revising the work to the final format.

## Footnotes

1 Sequence (seq)-1: CFLGYFCTCYFGLFC, seq-2: CFLGYCCCCYFGLFC, seq-3: CYLIVGYFCTCYFGVF, seq-4: CYLVVGYFCTCYFGVF, seq-5: CYLLVGYLCTCYFGVF, seq-6: FIGYVCTCYFGVF, seq-7: GYFCTVYFGVF, seq-8: LGFMCTCYFGVF, seq-9: GWFCTCYFGLY.

## References

1 Kerkar N, Vergani D. De novo autoimmune hepatitis –is this different in adults compared to children. J Autoimmun. 2018;26–33:26–33.

2 Metlas R, Skerl V, Veljkovic V, Colombatti A, Pongor S. Immunoglobulin-like domain of HIV-1 envelope glycoprotein gpl20 encodes putative internal image of some common human proteins. Viral Immunol. 1994;7(4):215–219.

3 Veljkovic V, Johnson E, Metlaš R. Molecular basis of the inefficacy and possible harmful effects of AIDS vaccine candidates based on HIV-1 envelope glycoprotein gp120. Vaccine. 1997;15(2):473–474.

4 Metlas R, Trajkovic D, Srdic T, Veljkovic V, Colombatti A. Human immunodeficiency virus V3 peptide-reactive antibodies are present in normal HIV-negative sera. AIDS Res Hum Retrovir. 1999;15(7):671–677.

5 Metlas R, Trajkovic D, Srdic T, Veljkovic V, Colombatti A. Anti-V3 and anti-IgG antibodies of healthy individuals share complementarity structures. J Acquir Immune Defic Syndr. 1999;21(4):266–270.

6 Carter CJ. Extensive viral mimicry of 22 AIDS-related autoantigens by HIV-1 proteins and pathway analysis of 561 viral/human homologues suggest an initial treatable autoimmune component of AIDS. FEMS Immunol Med Microbiol. 2011;63:254–268.

7 Kanduc D, Shoenfeld Y. From HBV to HPV: Designing vaccines for extensive and intensive vaccination campaigns worldwide. Autoimmun Rev. 2016;15:1054–1061.

8 Kanduc D, Shoenfeld Y. Inter-pathogen peptide sharing and the original antigenic sin: solving a paradox. The Open Immunology Journal. 2018;8:16–27.

9 Kanduc D, Shoenfeld Y. Human Papillomavirus epitope eimicry and autoimmunity: the molecular truth of peptide sharing. Pathobiology. 2019;86:285–295.

10 Kanduc D, Shoenfeld Y. On the molecular determinants of the SARS-CoV-2 attack. Clin Immunol. 2020;215:108426.

11 Kanduc D, Shoenfeld Y. Medical, genomic, and evolutionary aspects of the peptide sharing between pathogens, primates, and humans. Global Med Genet. 2020;7:64–67.

12 Kanduc D, Shoenfeld Y. Molecular mimicry between SARS-CoV-2 spike glycoprotein and mammalian proteomes: implications for the vaccine. Immunol Res. 2020;68:310–313.

13 Atyeo C, Fischinger S, Zohar T, Slein MD, Burke J, Loos C, McCulloch DJ, Newman KL, Wolf C, Yu J, et al. Distinct early serological signatures track with SARS-CoV-2 survival. Immunity. 2020;53:524–532.

14 Kaneko N, Kuo HH, Boucau J, Farmer JR, Allard-Chamard H, Mahajan VS, Piechocka-Trocha A, Lefteri K, Osborn M, Bals J, et al. Loss of Bcl-6-expressing T follicular helper cells and germinal centers in COVID-19. Cell. 2020;183:1–15.

15 Kuri-Cervantes L, Pampena MB, Meng W, Rosenfeld AM, Ittner CA, Weisman AR, Agyekum RS, Mathew D, Baxter AE, Vella LA, et al. Comprehensive mapping of immune perturbations associated with severe COVID-19. Sci Immunol. 2020;5:eabd7114.

16 Laing AG, Lorenc A, del Molino del Barrio I, Das A, Fish M, Monin L, Muñoz-Ruiz M, McKenzie DR, Hayday TS, Francos-Quijorna I, et al. A dynamic COVID-19 immune signature includes associations with poor prognosis. Nat Med. 2020;26:1623–1635.

17 Lucas C, Wong P, Klein J, Castro TB, Silva J, Sundaram M, Ellingson MK, Mao T, Oh JE, Israelow B, et al. Longitudinal analyses reveal immunological misfiring in severe COVID-19. Nature. 2020;584:463–469.

18 Mathew D, Giles JR, Baxter AE, Oldridge DA, Greenplate AR, Wu JE, Alanio C, Kuri-Cervantes L, Pampena MB, D’Andrea K, et al. Deep immune profiling of COVID-19 patients reveals distinct immunotypes with therapeutic implications. Science. 2020;369:eabc8511.

19 Woodruff MC, Ramonell RP, Cashman KS, Nguyen DC, Saini AS, Haddad N, Ley AM, Kyu S, Howell JC, Ozturk T, et al. Dominant extrafollicular B cell responses in severe COVID-19 disease correlate with robust viral-specific antibody production but poor clinical outcomes. [Internet].MedRxiv; 2020.

20 Bastard P, Rosen LB, Zhang Q, Michailidis E, Hoffmann HH, Zhang Y, Dorgham K, Philippot Q, Rosain J, Beziat V, et al. Autoantibodies against type I IFNs in patients with life-threatening COVID-19. Science. 2020;370:eabd4585.

21 Rodríguez Y, Novelli L, Rojas M, De Santis M, Acosta-Ampudia Y, Monsalve DM, Ramírez-Santana C, Costanzo A, Ridgway WM, Ansari AA, et al. Autoinflammatory and autoimmune conditions at the crossroad of COVID-19. J Autoimmun. 2020;114:102506.

22 Dotan A, Muller S, Kanduc D, David P, Halpert G, Shoenfeld Y. The SARS-CoV-2 as an instrumental trigger of autoimmunity. Autoimmun Rev. 2021 102792.

23 Cappello F, Gammazza AM, Dieli F, de Macario EC, Macario AJ. Does SARS-CoV-2 trigger stress-induced autoimmunity by molecular mimicry? A hypothesis. J Clin Med. 2020;9:2038.

24 Lucchese G, Flöel A. Molecular mimicry between SARS-CoV-2 and respiratory pacemaker neurons. Autoimmun Rev. 2020;19:102556.

25 Lucchese G, Flöel A. SARS-CoV-2 and Guillain-Barré syndrome: molecular mimicry with human heat shock proteins as potential pathogenic mechanism. Cell Stress Chaperones. 2020;25:731–735.

26 Angileri F, Legare S, Gammazza AM, de Macario EC, Macario AJ, Cappello F. Molecular mimicry may explain multi-organ damage in COVID-19. Autoimmun Rev. 2020;19:102591.

27 Vojdani A, Kharrazian D. Potential antigenic cross-reactivity between SARS-CoV-2 and human tissue with a possible link to an increase in autoimmune diseases. Clin Immunol. 2020;217:108480.

28 Adiguzel Y. Peptides of H. sapiens and P. falciparum that are predicted to bind strongly to HLA-A*24:02 and homologous to a SARS-CoV-2 peptide. [Internet].ArXiv; 2021.

29 Blanquart S, Gascuel O. Mitochondrial genes support a common origin of rodent malaria parasites and Plasmodium falciparum’s relatives infecting great apes. BMC Evol Biol. 2011;11:70.

30 Déchamps S, Maynadier M, Wein S, Gannoun-Zaki L, Maréchal E, Vial HJ. Rodent and nonrodent malaria parasites differ in their phospholipid metabolic pathways. J Lipid Res. 2010;51:81–96.

31 Altschul SF, Madden TL, Schäffer AA, Zhang J, Zhang Z, Miller W, Lipman DJ. Gapped BLAST and PSI-BLAST: a new generation of protein database search programs. Nucleic Acids Res. 1997;25:3389–3402.

32 NCBI Resource Coordinators. Database resources of the National Center for Biotechnology Information. Nucleic Acids Res. 2017;46:D8–D13.

33 Nguyen A, David JK, Maden SK, Wood MA, Weeder BR, Nellore A, Thompson RF. Human Leukocyte Antigen Susceptibility Map for Severe Acute Respiratory Syndrome Coronavirus 2. J Virol. 2020;94(13):e00510–20.

34 Andreatta M, Nielsen M. Gapped sequence alignment using artificial neural networks: application to the MHC class I system. Bioinformatics. 2016;32(4):511–517.

35 Nielsen M, Lundegaard C, Worning P, Lauemoller SL, Lamberth K, Buus S, Brunak S, Lund O. Reliable prediction of T-cell epitopes using neural networks with novel sequence representations. Protein Sci. 2003;12:1007–1017.

36 Reynisson B, Alvarez B, Paul S, Peters B, Nielsen M. NetMHCpan-4.1 and NetMHCIIpan-4.0: improved predictions of MHC antigen presentation by concurrent motif deconvolution and integration of MS MHC eluted ligand data. Nucleic Acids Res. 2020;48(W1):W449–W454.

37 Zhang H, Lund O, Nielsen M. The PickPocket method for predicting binding specificities for receptors based on receptor pocket similarities: application to MHC-peptide binding. Bioinformatics. 2009;25(10):1293–1299.

38 Karosiene E, Lundegaard C, Lund O, Nielsen M. NetMHCcons: a consensus method for the major histocompatibility complex class I predictions. Immunogenetics. 2012;64(3):177–186.

39 Stranzl T, Larsen MV, Lundegaard C, Nielsen M. NetCTLpan. Pan-specific MHC class I pathway epitope predictions. Immunogenetics. 2020;62(6):357–368.

40 The UniProt Consortium. UniProt: a worldwide hub of protein knowledge. Nucleic Acids Res. 2019;47(D1):D506–D515.

41 Stelzer G, Rosen R, Plaschkes I, Zimmerman S, Twik M, Fishilevich S, Stein TI, Nudel R, Lieder I, Mazor Y, et al. The GeneCards suite: from gene data mining to disease genome sequence analysis. Curr Protoc Bioinformatics. 2016;54:1.30.1–1.30.33.

42 Waterhouse AM, Procter JB, Martin DM, Clamp M, Barton GJ. Jalview version 2 – a multiple sequence alignment editor and analysis workbench. Bioinformatics. 2009;25(9):1189–1191.

43 Bianchi F, Textor J, van den Bogaart G. Transmembrane helices are an overlooked source of Major Histocompatibility Complex Class I epitopes. Front Immunol. 2017;8:1118.

44 Adiguzel Y. Molecular mimicry between SARS-CoV-2 and human proteins. Autoimmun Rev. 2021;20:102791.

45 Lyons-Weiler J. Pathogenic priming likely contributes to serious and critical illness and mortality in COVID-19 via autoimmunity. Journal of Translational Autoimmunity. 2020;3:100051.

46 Kanduc D. From anti-SARS-CoV-2 immune responses to COVID-19 via molecular mimicry. Antibodies. 2020;9:33.

47 Lucchese G. Cerebrospinal fluid findings in COVID-19 indicate autoimmunity. Lancet Microbe. 2020;1:e242.

48 Kohm AK, Fuller KG, Miller SD. Mimicking the way to autoimmunity: an evolving theory of sequence and structural homology. Trends Microbiol. 2003;11:101–105.

49 Lule S, Colpak AI, Balci-Peynircioglu B, Gursoy-Ozdemir Y, Peker S, Kalyoncu U, Can A, Tekin N, Demiralp D, Dalkara T. Behcet Disease serum is immunoreactive to neurofilament medium which share common epitopes to bacterial HSP-65, a putative trigger. J Autoimmun. 2017;84:87–96.

50 Negi S, Singh H, Mukhopadhyay A. Gut bacterial peptides with autoimmunity potential as environmental trigger for late onset complex diseases: in-silico study. PLoS One. 2017;12:e0180518.

51 Trost B, Lucchese G, Stufano A, Bickis M, Kusalik A, Kanduc D. No human protein is exempt from bacterial motifs, not even one. Self Nonself. 2010;1:328–334.

52 Vellozzi C, Iqbal S, Broder K. Guillain-Barre syndrome, influenza, and influenza vaccination: the epidemiologic evidence. Clin Infect Dis. 2014;58:1149–1155.

53 Yuki N. Ganglioside mimicry and peripheral nerve disease. Muscle Nerve. 2007;35:691–711.

54 Zabriskie JB, Freimer EH. An immunological relationship between the group. A streptococcus and mammalian muscle. J Exp Med. 1966;124:661–678.

55 Amela I, Cedano J, Querol E. Pathogen proteins eliciting antibodies do not share epitopes with host proteins: a bioinformatics approach. PLoS One. 2007;2(6):e512.

56 Matzinger P. The danger model: a renewed sense of self. Science. 2002;296:301–305.

57 Rojas M, Restrepo-Jiménez P, Monsalve DM, Pacheco Y, Acosta-Ampudia Y, Ramírez-Santana C, Leung PS, Ansari AA, Gershwin ME, Anaya JM. Molecular mimicry and autoimmunity. J Autoimmun. 2018;95:100–123.

58 Fujinami RS, Oldstone MB, Wroblewska Z, Frankel ME, Koprowski H. Molecular mimicry in virus infection: crossreaction of measles virus phosphoprotein or of herpes simplex virus protein with human intermediate filaments. Proc Natl Acad Sci Unit States Am. 1983;80:2346–2350.

59 Fujinami RS, Oldstone MB. Amino acid homology between the encephalitogenic site of myelin basic protein and virus: mechanism for autoimmunity. Science. 1985;230(4729):1043–1045.

60 Kanduc D. The comparative biochemistry of viruses and humans: an evolutionary path towards autoimmunity. Biol Chem. 2019;400:629–638.

61 Ahmed SF, Quadeer AA, McKay MR. Preliminary identification of potential vaccine targets for the COVID-19 coronavirus (SARS-CoV-2) based on SARS-CoV immunological studies. Viruses. 2020;12:254.

62 Kanduc D. Peptide cross-reactivity: the original sin of vaccines. Front Biosci. 2012;4:1393–1401.

63 Robson B. Computers and viral diseases. Preliminary bioinformatics studies on the design of a synthetic vaccine and a preventative peptidomimetic antagonist against the SARS-CoV-2 (2019-nCoV, COVID-19) coronavirus. Comput Biol Med. 2020;119:103670.

64 De Maio F, Lo Cascio E, Babini G, Sali M, Longa SD, Tilocca B, Roncada P, Arcovito A, Sanguinetti M, Scambia G, et al. Improved binding of SARS-CoV-2 envelope protein to tight junction-associated PALS1 could play a key role in COVID-19 pathogenesis. Microb Infect. 2020;22(10):592–597.

65 Teoh KT, Siu YL, Chan WL, Schlüter MA, Liu CJ, Peiris JS, Bruzzone R, Margolis B, Nal B. The SARS Coronavirus E protein interacts with PALS1 and alters tight junction formation and epithelial morphogenesis. Mol Biol Cell. 2010;21(22):3838–3852.

66 Schoeman D, Fielding BC. Is there a link between the pathogenic human coronavirus envelope protein and immunopathology? A review of the literature. Front Microbiol. 2020;11:2086.

67 Lo Cascio E, Toto A, Babini G, De Maio F, Sanguinetti M, Mordente A, Longa SD, Arcovito A. Structural determinants driving the binding process between PDZ domain of wild type human PALS1 protein and SLiM sequences of SARS-CoV E proteins. Comput Struct Biotechnol J. 2021;19:1838–1847.

68 Ziegler CG, Miao VN, Owings AH, Navia AW, Tang Y, Bromley JD, Lotfy P, Sloan M, Laird H, Williams HB, et al. Impaired local intrinsic immunity to SARS-CoV-2 infection in severe COVID-19. [Internet].BioRxiv; 2021.

69 Rappaport N, Twik M, Plaschkes I, Nudel R, Stein TI, Levitt J, Gershoni M, Morrey CP, Safran M, Lancet D. MalaCards: an amalgamated human disease compendium with diverse clinical and genetic annotation and structured search. Nucleic Acids Res. 2017;45(D1):D877–D887.

70 Loukovitis E, Anastasia S, Tranos P, Balidis M, Asteriadis S, Thanos V, Thanos S, Anogeianakis G. A review of recent developments in retinitis pigmentosa genetics, its clinical features, and natural course. Med Hypothesis Discov Innov Ophthalmol. 2020;9(4):231–254.

71 Warren RL, Birol I. HLA Predictions from the bronchoalveolar lavage fluid samples of five patients at the early stage of the Wuhan seafood market COVID-19 outbreak. [Internet].ArXiv; 2020.

72 Cheranev V, Loginova M, Jankevic T, Kutyavina S, Rebrikov D. A novel allele, HLA-C*15:227, identified when typing COVID-19 patients. HLA. 2021;97(4):377–378.

73 Robinson J, Barker DJ, Georgiou X, Cooper MA, Flicek P, Marsh SG. IPD-IMGT/HLA database. Nucleic Acids Res. 2020;48(D1):D948–D955.

